# Histone Demethylase KDM6B Promotes Postnatal Oligodendrocyte Development and Cortical Myelination

**DOI:** 10.64898/2025.12.30.697092

**Authors:** Ruth Lambries, Zihan Shen, George I Mias, Jin He

**Author notes:** The authors contributed to the work equally (Ruth Lambries, Zihan Shen).

## Abstract

Myelination in the postnatal cortex requires epigenetic programs that activate oligodendrocyte gene networks. Here we show that the histone H3K27me3 demethylase KDM6B promotes this process *in vivo*. Conditional *Kdm6b* deletion (*Kdm6b*-cKO) in Emx1 dorsal telencephalic progenitors caused delayed cortical myelination. Lineage-specific RNA-seq revealed that the oligodendrocyte master regulator *Sox10* was significantly reduced, as well as myelination effectors, indicating impaired oligodendrocyte maturation. Chromatin immunoprecipitation assays of the same lineage showed increased H3K27me3 at promoter-proximal regions of *Sox10,* consistent with loss of KDM6B-mediated de-repression. Temporally controlled *Sox10* induction partially rescued myelination in *Kdm6b*-cKO mice. These findings define a KDM6B→SOX10 axis that licenses oligodendrocyte development and cortical myelination by removing repressive H3K27me3 and sustaining active chromatin, with implications for KDM6B-related neurodevelopmental disorders.

## Introduction

Myelination is a hallmark of vertebrate brain maturation, enabling rapid saltatory conduction and supporting metabolic and trophic coupling between axons and glia(Fletcher et al., 2021; Zhang et al., 2025). In the neocortex, oligodendrocyte lineage progression—from progenitors to myelinating oligodendrocytes—unfolds predominantly postnatally and is tightly coordinated with circuit assembly and activity(Kuhn et al., 2019). Transcriptional regulators such as SOX10 orchestrate this program by driving expression of myelin structural genes (e.g., *MBP*) and lipid biosynthetic pathways required for sheath formation and stability(Pozniak et al., 2010; Elbaz and Popko, 2019). While these lineage determinants are well characterized, the epigenetic mechanisms that poise and activate the myelin gene network in the postnatal cortex remain poorly understood(Liu et al., 2016).

A major layer of epigenetic control involves covalent histone modifications that shape chromatin accessibility and gene expression. Trimethylation of histone H3 lysine 27 (H3K27me3)—deposited by the Polycomb Repressive Complex 2 (PRC2)—is a canonical repressive mark that constrains developmental competence(Cao et al., 2002; Mozgova et al., 2015). The histone demethylase KDM6B catalyzes removal of H3K27me3, thereby antagonizing PRC2 and facilitating transcriptional activation at developmental loci(Agger et al., 2007; Sen et al., 2008). Beyond its biochemical activity, KDM6B has been implicated in numerous neural processes, including neural commitment from pluripotent stem cells, neuronal and glial lineage specification, activity-dependent neuronal survival, postnatal neurogenesis, and neuronal maturation(Burgold et al., 2008; Park et al., 2014; Wijayatunge et al., 2014; Wijayatunge et al., 2018; Shan et al., 2020). These observations raise the possibility that KDM6B might also be required for postnatal oligodendrocyte development and cortical myelination, where timely release from Polycomb repression is likely essential.

The clinical relevance of KDM6B is underscored by human genetics. Recent large-scale studies identify KDM6B as a high-confidence autism spectrum disorder (ASD) risk gene, and multiple case reports describe children with disruptive or missense KDM6B variants presenting with neurodevelopmental phenotypes that include intellectual disability, delayed language/motor development, and attention-deficit/hyperactivity disorder (ADHD) behaviors(Stolerman et al., 2019; Satterstrom et al., 2020; Insa Pineda and Gomez Gonzalez, 2021; Rots et al., 2023, 2025). Together with experimental evidence that heterozygous loss of *Kdm6b* in mice produces ASD-and ADHD-like behaviors, these findings suggest that KDM6B-dependent chromatin regulation contributes to brain development and function(Gao et al., 2022). Whether defects in oligodendrocyte maturation and myelination participate in the pathophysiology of KDM6B-related neurodevelopmental disorders, however, remains an open question.

Here we investigate the function of KDM6B in licensing the oligodendrocyte/myelin program in the developing cortex through removal of H3K27me3 at key regulatory genes. We conditionally deleted *Kdm6b* in Emx1-positive dorsal telencephalic progenitors, enabling us to probe KDM6B function within a lineage that contributes to cortical glutamatergic neurons, astrocytes, and a subset of oligodendrocytes. To define consequences at tissue and molecular levels, we combined postnatal myelin histology, lineage-specific RNA-seq, and chromatin immunoprecipitation (ChIP) assays for H3K27me3. Finally, we examined causality by temporal, Tet-inducible *Sox10* overexpression *in vivo* as a targeted rescue of a predicted KDM6B effector.

Our results show that loss of *Kdm6b* delays postnatal cortical myelination, coincident with downregulation of *Sox10* early and subsequent reduction of myelination effector genes by P30. At the chromatin level, *Kdm6b***-**deficient Emx1-lineage cells exhibit increased H3K27me3 at promoter-proximal regions of *Sox10*, consistent with loss of KDM6B-mediated de-repression. Importantly, induced *Sox10* expression in the developing cortex partially rescues myelination in *Kdm6b*-cKO mice, establishing SOX10 as a functionally significant downstream effector of KDM6B in this context.

Collectively, these findings define a KDM6B→SOX10 axis that promotes oligodendrocyte maturation and cortical myelination, illuminate how histone demethylation interfaces with lineage transcription factors during postnatal brain development, and provide mechanistic traction for understanding—and potentially treating—KDM6B-related neurodevelopmental disorders.

## Materials and Methods

### Mice

Founder conditional *Kdm6b* knockout (*Kdm6b*-cKO) mice were generated by *in vitro* fertilization at the Michigan State University (MSU) Transgenic Core, as reported in our previous study(Gao et al., 2022). Mice were housed under standard conditions (12 h light/12 h dark cycle) with ad libitum access to food and water. All procedures were approved by the Michigan State University Institutional Animal Care and Use Committee (IACUC, PROTO202300162) and conducted in accordance with institutional guidelines.

### Mouse breeding strategy

All mice were backcrossed to C57BL/6 for more than five generations to achieve a congenic C57BL/6 background. Conditional knockouts and controls were generated by crossing Emx1-Cre mice (B6.129S2-Emx1^tm1^(cre)^Krj^/J, The Jackson Laboratory) with *Kdm6b* floxed mice. Littermates included wild-type (*Kdm6b*^wt/wt^) and conditional knockout (Emx1-Cre; Kdm6b^f/f^) animals. For breeding, *Kdm6b*^wt/f^ × *Kdm6b*^wt/f^ matings produced *Kdm6b*^wt/wt^, *Kdm6b*^wt/f,^ and *Kdm6b*^f/f^ offspring. All lines were maintained on a SUN1-GFP reporter background (B6;129-*Gt(ROSA)26Sor^tm5(CAG-Sun1/sfGFP)Nat^*/J, The Jackson Laboratory).

### In uterus electroporation

E15.5 dams were anesthetized with isoflurane on a 37 °C heated stage, the abdomen was aseptically prepared, and a midline laparotomy was performed to externalize the uterine horns on warm, saline-moistened gauze. Endotoxin-free plasmids (1 µg/µL total in sterile PBS with 0.01% Fast Green) were loaded into a pulled glass capillary attached to a mouth pipette and 1.0 µL DNA was injected into the lateral ventricle. Tweezer-type platinum electrodes were positioned to direct the anode toward the dorsal pallium and 5 square-wave pulses were delivered with a pulse generator (40 V, 50 ms width, 950 ms interval). After electroporation, uterine horns were returned to the abdominal cavity, the peritoneum and skin were closed with absorbable sutures and wound clips, and dams received carprofen per IACUC guidelines. The procedure was approved by the Michigan State University Institutional Animal Care and Use Committee (IACUC) and conducted in accordance with institutional guidelines.

### Doxycycline administration

To induce gene expression in offspring carrying a doxycycline-inducible transgene, pregnant dams were provided with doxycycline-containing drinking water starting at embryonic day 18.5 (E18.5) and maintained continuously until postnatal day 21 (P16). The doxycycline solution was prepared fresh every 2 days by dissolving doxycycline hyclate (Sigma-Aldrich) at a final concentration of 2 mg/mL in 2% sucrose (w/v) to improve palatability and protected from light using amber bottles. Dams were housed under standard conditions with ad libitum access to the doxycycline water, which ensured sustained systemic exposure through lactation to activate the tetracycline-responsive transgene in the pups during late embryonic and early postnatal development. The procedure was approved by the Michigan State University Institutional Animal Care and Use Committee (IACUC) and conducted in accordance with institutional guidelines

### Tissue processing for histology

P16 and P21 mice were anesthetized by avertin and transcardially perfused with ice-cold phosphate-buffered saline followed by 4% paraformaldehyde (PFA) in PBS. Brains were dissected and post-fixed in 4% PFA at 4 °C overnight, then cryoprotected in 30% sucrose in PBS at 4 °C until tissue sank. Cryoprotected brains were embedded in optimal cutting temperature (OCT) compound in molds, frozen on dry-ice or a pre-chilled isopentane bath, and stored at −80 °C. Coronal sections were cut at 10 µm on a cryostat (Leica CM1950).

### Immunostaining

Slide-mounted OCT cryosections were rinsed in PBS and permeabilized/blocked for 1 h at RT in PBS containing 0.3% Triton X-100 and 5% normal donkey. Rabbit anti-MBP (Cell Signaling) and Rabbit anti-SOX10 (Cell Signaling) diluted in the same buffer (1:400) were applied overnight at 4 °C in a humidified chamber. The next day, sections were washed in PBS and incubated with Donkey anti-rabbit Alexa Fluor 680–conjugated secondary antibodies (1:400, Jackson ImmunoResearch) for 1 h at RT protected from light, followed by PBS washes. Nuclei were counterstained with DAPI (0.5 µg/mL, 5 min), briefly rinsed, and coverslipped with antifade mounting medium.

### Imaging

Images were acquired using a ZEISS Axio Observer Z1 inverted fluorescence microscope equipped with 2.5× and 10× objective lenses and captured with ZEN Pro software (ZEISS).

### Imaging data analysis

Mean fluorescence intensity (MFI) of MBP immunofluorescence was quantified using ImageJ (NIH). All images were converted to 8-bit format to standardize intensity scaling across samples. One image per sample was analyzed. A region of interest (ROI) encompassing a high-density myelinated area immediately above the corpus callosum was used as a reference region and set as 100 for normalization. The sampling ROI encompassed the outer extent of cortical myelinated axons and was used as the primary measurement region for MFI quantification. For the myelinated fiber length proportion analysis, the same sections used for MFI analysis were used to quantify cortical myelination length using ImageJ. Total cortical width was measured by obtaining five independent linear measurements (in pixels) from the top of the corpus callosum to the pial surface within the superior lateral cortex. These measurements were averaged to yield a mean cortical width for each sample. The extent of myelination was determined by taking five measurements from the top of the corpus callosum to the distal boundary of the myelinated region. These values were averaged to obtain the mean myelinated length. The proportion of cortical myelination was calculated by dividing the mean myelinated length by the mean cortical width for each sample. This analysis was performed for all samples used in the study. SOX10 immunostaining was quantified using ImageJ. All images were converted to 8-bit format to allow automated particle analysis. For each sample, three 400 × 400-pixel ROIs were randomly selected within the analyzed hemisphere. Automated particle counting was performed for each ROI using ImageJ’s particle analysis function. Cellular density was calculated by dividing the particle count by the ROI area (160,000 pixels²). Density values from the three ROIs were averaged to obtain a single value per sample for statistical analysis.

### Fluorescence-activated nuclei sorting

Mouse cortices were rapidly dissected on ice. Nuclei were isolated sucrose gradient centrifugation. Briefly, 14 mL of sucrose cushion (1.8 M sucrose, 10 mM Tris-HCl pH 8.0, 3 mM MgCl ) was added to the bottom of Beckman Coulter ultracentrifuge tubes. Using a glass Dounce homogenizer, a freshly isolated or frozen cortex was homogenized in 12 mL homogenization buffer (0.32 M sucrose, 5 mM CaCl 3 mM MgCl , 10 mM Tris-HCl pH 8.0, 0.1% Triton X-100, 0.1 mM EDTA with strokes of the loose pestle followed by strokes of the tight pestle. Homogenates (∼12 mL) were carefully layered onto the sucrose cushion, and 10 mL homogenization buffer was added on top. Gradients were centrifuged at 25,000 rpm for 2.5 h at 4 °C in a Beckman Coulter L8-70M ultracentrifuge using an SW28 swinging-bucket rotor. Supernatants were removed by aspiration. Nuclei were resuspended in 0.01% BSA in DPBS passed through a 40-µm cell strainer. The SUN1-GFP+ nuclei were sorted by S3e sorter (Bio-Rad).

### qRT-PCR assays

Total RNA was extracted using QIAshredder and RNeasy spin columns (QIAGEN) according to the manufacturer’s instructions. RNA was reverse-transcribed with HIScript Reverse Transcriptase III (Vazyme). Quantitative PCR was performed using SupRealQ Ultra Hunter SYBR qPCR Master Mix (U+) (Vazyme) on a CFX384 Touch Real-Time PCR Detection System (Bio-Rad). Primer sequences are listed in Supplementary Table 1.

### RNA-seq sample preparation for AVITI sequencing

Total RNA was extracted using per the manufacturer’s instructions. RNA-seq libraries were repared from total RNA using the VAHTS Universal V10 RNA-seq Library Prep Kit for Illumina (Vazyme). Adapter-ligated cDNA was PCR-amplified and size-selected magnetic beads. DNA was measured by an Invitrogen Qubit fluorometer. Libraries were sequenced on NovaSeq6000 or Element AVITI sequencers at the Michigan State University Genomics Core.

### RNA-Seq data analysis

RNA-seq analysis was performed essentially as described previously(Aljazi et al., 2021; Gao et al., 2021). Reads were aligned to the mouse reference genome (UCSC mm9) using HISAT2(Kim et al., 2019). Gene-level abundance was normalized as reads per kilobase per million mapped reads (RPKM). Differential expression was assessed with Cuffdiff(Trapnell et al., 2012), using a false-discovery rate threshold of q < 0.05 and a > 2-fold change in RPKM between conditions. Differentially expressed gene sets were submitted to g:Profiler (https://biit.cs.ut.ee/gprofiler/gost) for Gene Ontology enrichment analysis.

### ChIP-qPCR analysis

ChIP was performed as described previously(He et al., 2013). FANS-sorted nuclei (2–3 × 10^5^ per IP) were lysed in 1% SDS, 10 mM EDTA, 50 mM Tris-HCl (pH 8.0) supplemented with protease inhibitors. Chromatin was sonicated on a Covaris S220 to an average fragment size of ∼200–400 bp, clarified by centrifugation, and diluted 10× with ChIP dilution buffer (0.01% SDS, 1.1% Triton X-100, 1.2 mM EDTA, 16.7 mM Tris-HCl pH 8.0, 167 mM NaCl). Supernatants were incubated overnight at 4 °C with 2 µL rabbit polyclonal anti-H3K27me3 (Cell Signaling Technology; catalog and lot as in Antibody Table). Immune complexes were captured with Protein G Plus/Protein A magnetic beads (EMD Millipore). Beads were washed sequentially in low-salt, high-salt, LiCl, and TE buffers, eluted in 1% SDS/0.1 M NaHCO , and crosslinks were reversed (65 °C, ≥4 h). DNA was purified by phenol–chloroform extraction and ethanol precipitation and used for library preparation. Quantitative PCR was performed using SupRealQ Ultra Hunter SYBR qPCR Master Mix (U+) (Vazyme) on a CFX384 Touch Real-Time PCR Detection System (Bio-Rad). Primer sequences are listed in Supplementary Table 1.

### Statistical analysis

All statistical analyses were performed using GraphPad Prism 8 (GraphPad Software). For comparisons between two groups, statistical significance was assessed using two-tailed unpaired Student’s *t* tests. For comparisons involving multiple samples, group multiple-row *t* tests were applied with Holm–Šidák correction for multiple comparisons. *P* values < 0.05 were considered statistically significant. Data are presented as mean ± SEM.

## Results

### Loss of *Kdm6b* delays the postnatal cortical myelination

To examine the role of KDM6B in cortical neural development, we conditionally deleted *Kdm6b* in Emx1***-***positive dorsal telencephalic progenitors—which give rise to cortical glutamatergic neurons, astrocytes, and oligodendrocytes—by crossing *Kdm6b*^flox/flox^ mice with the Emx1-Cre line(Gorski et al., 2002). Both wild-type (WT) and *Kdm6b***-**cKO cohorts were crossed to a SUN1-GFP reporter line to label Emx1D progeny, enabling lineage tracing and FANS (Fluorescence-Activated Nuclei Sorting)-based isolation of Emx1-lineage cells from cortices for downstream characterization (Figure 1A). To confirm knockout efficiency, we sorted SUN1-GFP cortical nuclei and performed quantitative real-time PCR (qRT-PCR) to measure *Kdm6b* mRNA. Relative to wild type, *Kdm6b***-**cKO samples showed a **>**95% reduction in *Kdm6b* transcript levels (i.e., **<** 5% of WT), confirming efficient gene deletion (Figure 1B, C).

**Figure 1.**
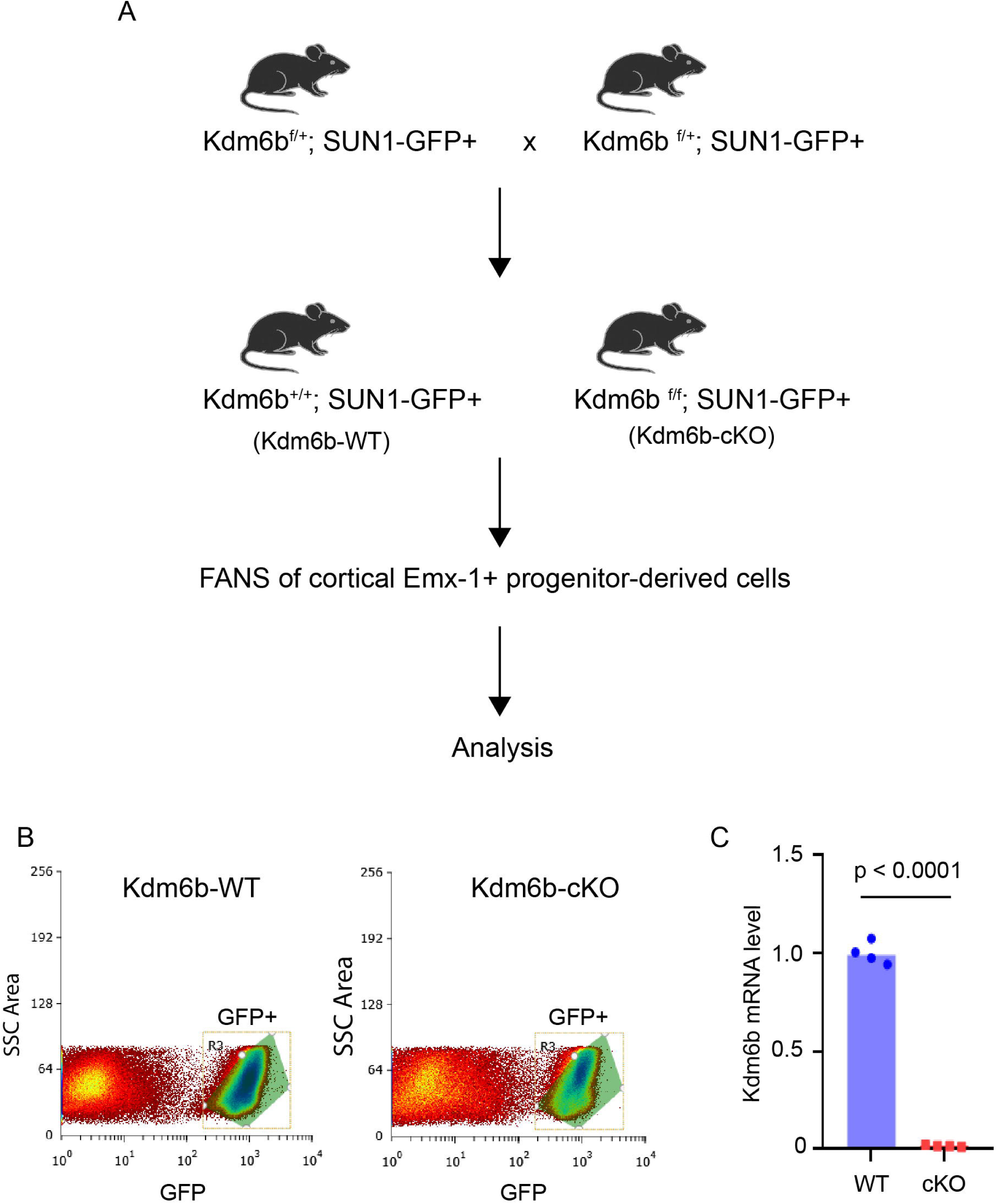
Generation and validation of *Kdm6b* conditional knockout in Emx1-lineage cortical cells. **(A)** Breeding scheme used to generate *Kdm6b*^+/+^; SUN1-GFP^+^ (WT) and Kdm6b^f/f^; SUN1-GFP^+^ (cKO) mice. Emx1-lineage cortical nuclei were isolated by fluorescence-activated nuclei sorting (FANS) based on GFP expression for downstream analyses. **(B)** Representative FANS plots showing gating of GFP^+^ cortical nuclei from WT and cKO mice. **(C)** qRT-PCR analysis of GFP^+^ nuclei confirming efficient loss of *Kdm6b* mRNA in cKO samples relative to WT controls. *Kdm6b* expression was normalized to *Gapdh*. Each dot represents one biological replicate.

To assess postnatal cortical myelination, we performed immunostaining for myelin basic protein (MBP) at postnatal day (P) 3, P16, and P24. As expected, wild-type (WT) mice showed a progressive increase in MBP signal—minimal/absent at P3, moderate at P16, and robust at P24. *Kdm6b*-cKO littermates followed a similar temporal trajectory but displayed markedly reduced MBP-positive neural fiber length and immunofluorescence intensity at P16 and P21 compared with WT (Figure 2), indicating that loss of *Kdm6b* in cortical neural progenitors impairs oligodendrocyte development and delays postnatal cortical myelination.

**Figure 2.**
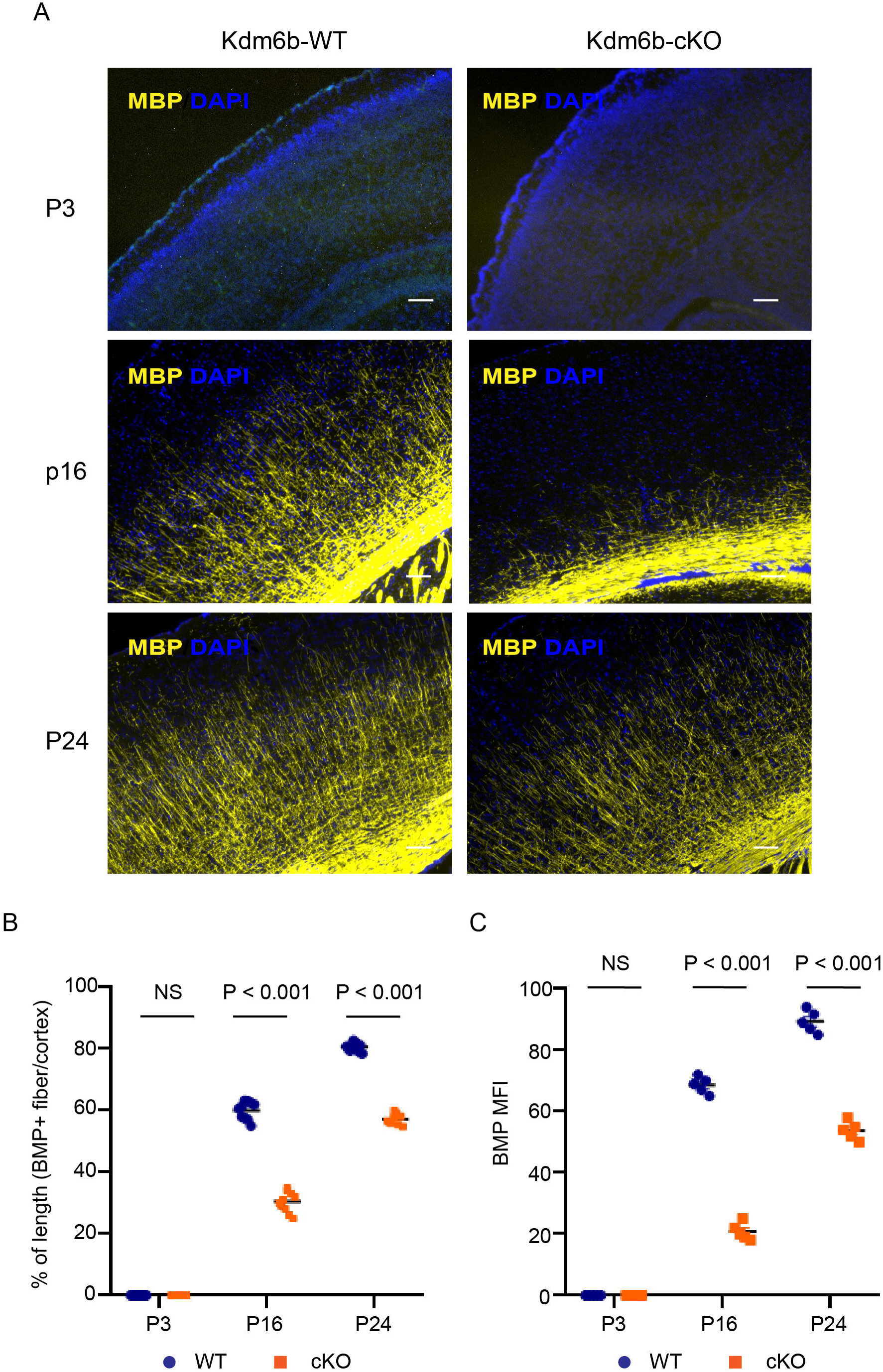
Delayed cortical myelination following *Kdm6b* deletion in Emx1-lineage cells. **(A)** Representative immunofluorescence images of cortical sections from *Kdm6b*-WT and *Kdm6b-cKO* mice at postnatal day 3 (P3), P16, and P24, stained for myelin basic protein (MBP, yellow) and DAPI (blue). WT cortices show robust, age-dependent increases in MBP-positive myelinated fibers, whereas *Kdm6b*-cKO cortices exhibit markedly reduced MBP labeling at P16 and P24. Scale bars, 200 μm. **(B)** Quantification of MBP-positive fiber length normalized to cortical length at P3, P16, and P24. No significant difference is observed at P3, whereas MBP-positive fiber length is significantly reduced in *Kdm6b*-cKO cortices at P16 and P24. Each dot represents an individual animal. Statistical significance is indicated (NS, not significant; *P* < 0.001).

### Loss of *Kdm6b* impairs expression of genes essential for postnatal oligodendrocyte development and function

To define the molecular mechanisms by which KDM6B regulates oligodendrocyte development and myelination, we performed RNA sequencing of Emx1 lineage–derived cortical cells isolated at postnatal day 1 (P1) and postnatal day 30 (P30). For both WT and *Kdm6b*-cKO littermates, cortical nuclei were purified by FANS using the SUN1-GFP reporter prior to library preparation. At P1, differential expression analysis identified approximately 122 downregulated and 327 upregulated genes in *Kdm6b*-cKO cortices relative to WT controls (fold changes > 2, q < 0.05). Notably, *Sox10*, a master regulator of oligodendrocyte lineage progression, was among the most significantly downregulated genes (Figure 3A, B). Consistent with the RNA-seq results, SOX10 immunostaining of P1 cortices revealed a marked reduction in SOX10 protein levels in *Kdm6b*-cKO mice (Figure 3C, D).

**Figure 3.**
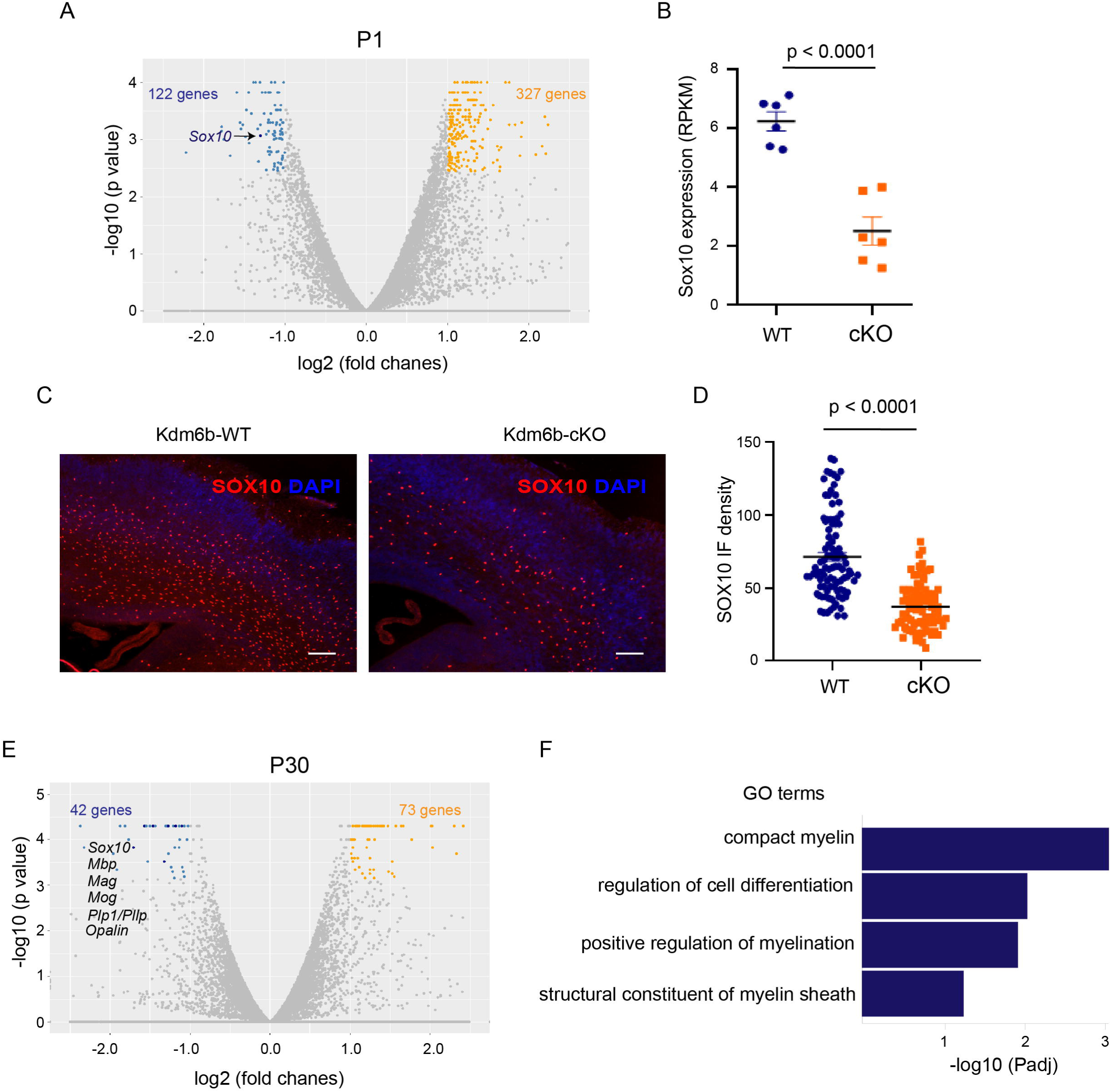
*Kdm6b* deletion suppresses *Sox10* expression and oligodendrocyte gene programs during cortical development. **(A)** Volcano plot of differentially expressed genes in P1 *Kdm6b*-cKO versus WT Emx1-lineage cortical cells. Blue and orange dots denote significantly downregulated and upregulated genes, respectively (|log FC| > 1, FDR < 0.05). *Sox10* is highlighted among the downregulated genes. **(B)** RNA-seq quantification showing reduced *Sox10* expression in *Kdm6b*-cKO cortices at P1, expressed as reads per kilobase per million mapped reads (RPKM). Data are mean ± SEM; ****p < 0.0001, two-tailed Student’s *t* test. **(C)** Representative immunofluorescence images of SOX10 (red) and DAPI (blue) in WT and *Kdm6b*-cKO cortices at P1. Scale bars, 200 μm. **(D)** Quantification of SOX10+ cell density in WT and *Kdm6b*-cKO cortices. Each dot represents an individual sampling region. Mean ± SEM; ****p < 0.0001. **(E)** Volcano plot of differentially expressed genes in P30 cortices, highlighting sustained downregulation of oligodendrocyte lineage genes including *Sox10, Mbp, Mag, Mog, Plp1,* and *Opalin* in *Kdm6b*-cKO mice. **(F)** Gene ontology (GO) enrichment analysis of downregulated genes in *Kdm6b*-cKO cortices at P30, revealing significant enrichment for myelin-related biological processes and oligodendrocyte differentiation programs.

By P30, transcriptomic analysis identified 42 downregulated and 73 upregulated genes. Notably, *Sox10* remained persistently suppressed, accompanied by broad downregulation of genes essential for oligodendrocyte maturation and myelin sheath formation, including *Mbp, Mag, Mog, Plp1/Pllp, Opalin,* and *Cmtm5* (Figure 3E). Gene ontology analysis further revealed significant enrichment of downregulated genes in myelin-associated biological processes, including compact myelin formation, structural constituents of the myelin sheath, and positive regulation of myelination (Figure 3F).

Together, these dynamic transcriptional changes closely parallel the histological deficits observed in *Kdm6b*-cKO cortices and indicate that impaired cortical myelination results from disrupted oligodendrocyte lineage progression driven by loss of key regulatory programs, most notably *Sox10*.

### Loss of *Kdm6b* alters promoter-associated histone H3K27me3 levels at the *Sox10* locus

To assess the epigenetic consequences of KDM6B loss, we performed chromatin immunoprecipitation followed by quantitative PCR (ChIP–qPCR) on FANS-sorted Emx1-lineage cortical cells, focusing on histone H3K27me3, a repressive modification deposited by Polycomb repressive complex 2 (PRC2) and removed by KDM6B. Compared with WT controls, *Kdm6b*-cKO cells exhibited significantly increased H3K27me3 enrichment at promoter-proximal regions of genes that were transcriptionally downregulated upon *Kdm6b* deletion. Notably, H3K27me3 levels were markedly elevated at the *Sox10* locus at amplicons near the transcription start site, whereas more distal regions showed no significant changes (Figure 4). These data indicate that KDM6B directly promotes gene activation during cortical development by erasing repressive H3K27me3 marks at gene promoters. Consistent with this mechanism, the reduced expression of *Sox10*, a master regulator of oligodendrocyte development, observed in *Kdm6b*-cKO cortices is likely a direct consequence of impaired promoter chromatin activation resulting from *Kdm6b* loss.

**Figure 4.**
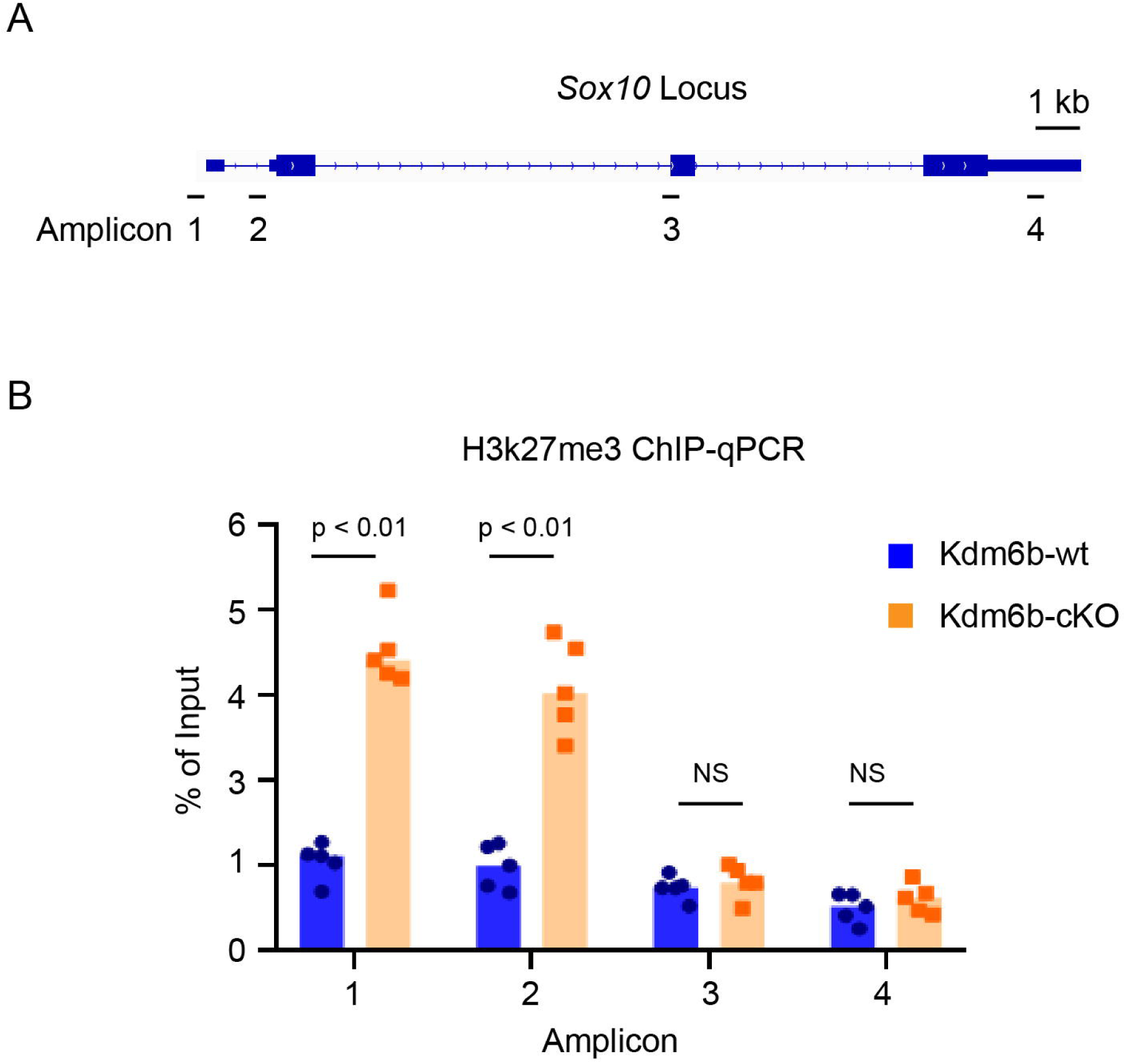
Loss of *Kdm6b* leads to increased H3K27me3 at the *Sox10* locus. **(A)** Schematic of the *Sox10* genomic locus showing exon–intron structure and the positions of qPCR amplicons (Amplicons 1–4) used for ChIP-qPCR analysis. **(B)** H3K27me3 ChIP-qPCR analysis at the *Sox10* locus in cortical tissue from *Kdm6b*-WT and *Kdm6b*-cKO mice. Enrichment is shown as percentage of input DNA. H3K27me3 levels are significantly increased at Amplicons 1 and 2 in *Kdm6b*-cKO samples compared with WT controls, whereas no significant differences are observed at Amplicons 3 and 4. Each dot represents one biological replicate; bars indicate mean ± SEM. Statistical significance is indicated (*p* < 0.01; ns, not significant).

### Ectopic expression of *Sox10* in the developing cortex partially rescues the cortical myelination

To test whether *Sox10* is a critical KDM6B target mediating postnatal oligodendrocyte development and cortical myelination, we attempted a rescue experiment by ectopically expressing *Sox10* in neural progenitors within the subventricular zone (SVZ) of the embryonic cortex using in utero electroporation (IUE) at E15.5. We initially employed a constitutive expression plasmid encoding Sox10–P2A–RFP, in which the self-cleaving P2A peptide generates separate SOX10 and RFP proteins, enabling fluorescent identification of transfected cells(Szymczak et al., 2004). A control plasmid expressing truncated LacZ–P2A–RFP was electroporated in parallel.

Constitutive *Sox10* overexpression at E15.5 profoundly disrupted radial migration, differentiation, and proliferation of cortical progenitors. Whereas LacZ-transfected neurons migrated normally and populated cortical layers II–III with typical neuronal morphology, Sox10-transfected progenitors largely failed to migrate and instead accumulated within the subventricular zone (SVZ). *Sox10*-expressing cells also exhibited reduced dispersion and prominent clustering near the ventricular region, consistent with impaired progenitor differentiation and proliferation (Figure S1). Moreover, these cells lacked neuronal characteristics, as indicated by the absence of RFP-labeled callosal axons. Together, these findings demonstrate that premature *Sox10* expression in the embryonic cortex diverts progenitors toward non-neuronal fates that are incompatible with normal neuronal migration, differentiation, and proliferation during cortical development.

To circumvent premature fate switching, we implemented a tetracycline-inducible system to achieve temporal control of *Sox10* expression(Gossen and Bujard, 1992). TRE–(Sox10–P2A–RFP) or TRE–(LacZ–P2A–RFP) responder plasmids were co-electroporated with an EF1α–rtTA activator. Validation in HEK293T cells demonstrated robust doxycycline (Dox)-dependent induction, with markedly increased RFP fluorescence upon Dox treatment and minimal basal expression in its absence. Immunoblotting further confirmed an approximately fivefold increase in SOX10 protein levels compared with controls (Figure S2).

Using this inducible system, Dox was administered from E18.5 to P16 to activate transgene expression in postmitotic or late-stage cells. Under these conditions, LacZ-electroporated cells predominantly migrated to cortical layers II–III, whereas *Sox10*-electroporated cells accumulated mainly in layers IV–VI, with fewer cells reaching superficial layers (Figure S3). Notably, inducible *Sox10* expression significantly increased cortical myelination in *Kdm6b*-cKO cortices at P16 compared with LacZ-transfected *Kdm6b*-cKO controls, although myelination did not fully reach wild-type levels (Figure 5). Together, these results indicate that *Sox10* overexpression partially rescues the myelination deficit caused by *Kdm6b* loss, supporting SOX10 as a key downstream effector of KDM6B in postnatal oligodendrocyte development.

**Figure 5.**
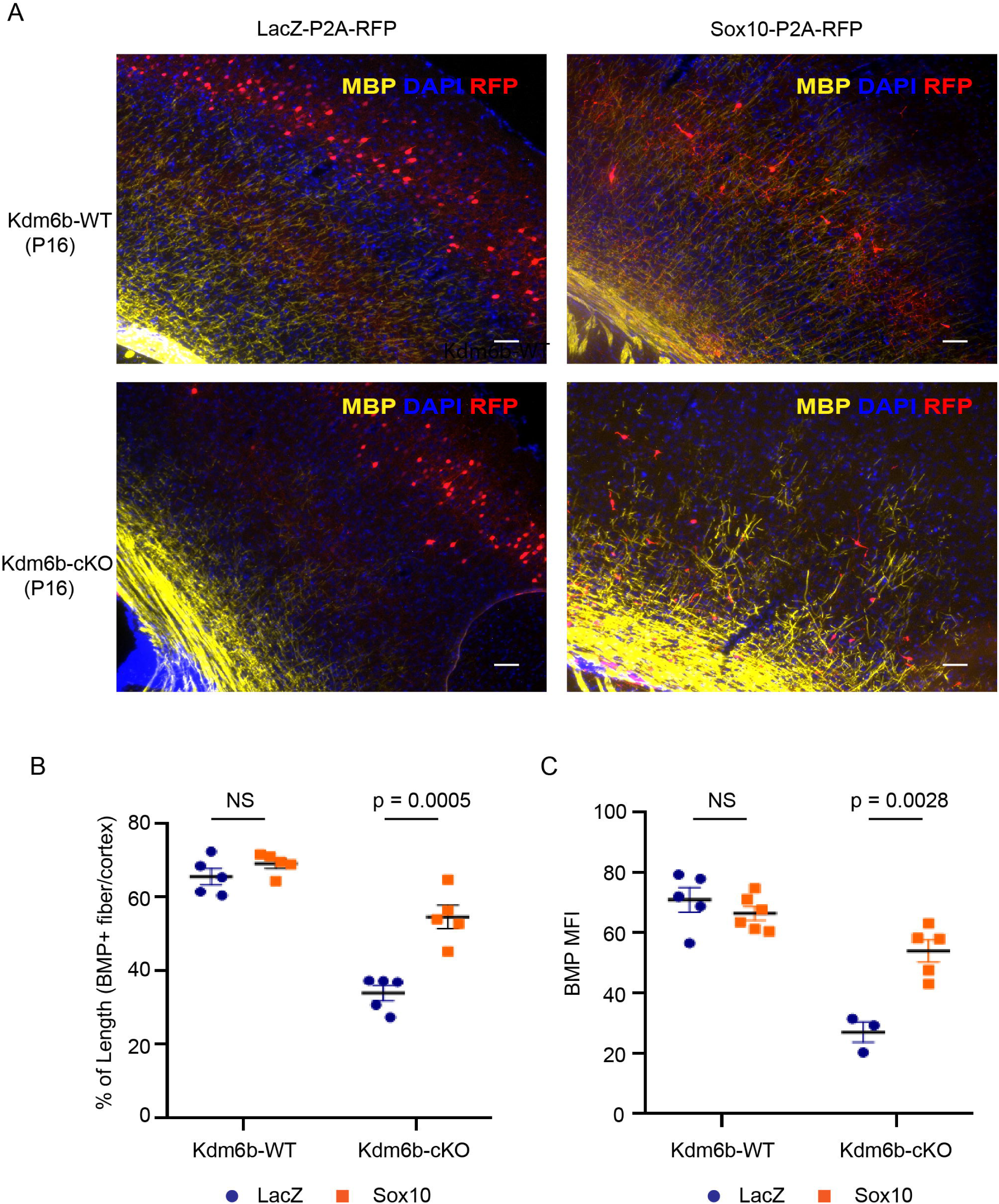
Inducible *Sox10* expression partially rescues cortical myelination in *Kdm6b*-cKO mice. **(A)** Representative immunofluorescence images of cortical sections from *Kdm6b*-WT and *Kdm6b*-cKO mice following in utero electroporation of LacZ–P2A–RFP (control) or Sox10–P2A–RFP constructs. Sections were stained for myelin basic protein (MBP, yellow), DAPI (blue), and RFP (red) to identify electroporated cells. **(B)** Quantification of MBP-positive fiber length normalized to total cortical length. *Sox10* expression does not significantly alter myelination in WT cortex but significantly increases MBP-positive fiber length in *Kdm6b*-cKO cortex compared with LacZ controls. **(C)** Quantification of mean MBP immunofluorescence intensity (MFI). *Sox10* overexpression significantly elevates MBP MFI in *Kdm6*b-cKO cortices, whereas no significant change is observed in WT cortex. Each dot represents one biological replicate; bars indicate mean ± SEM. Statistical significance is indicated (NS, not significant).

## Discussion

Our study identifies KDM6B as a pivotal epigenetic regulator of postnatal cortical myelination. Deletion of *Kdm6b* in Emx1 dorsal telencephalic progenitors resulted in a pronounced delay in cortical myelin accumulation during early postnatal development, with markedly reduced MBP immunoreactivity at P16–P21 (Figure 2). These anatomical deficits are mirrored at the molecular level: RNA-seq of Emx1-lineage cortical cells revealed downregulation of oligodendrocyte lineage regulators and myelin-associated genes, including early suppression of *Sox10* at P1 and reduced expression of myelination-related genes such as *Mbp* and *Pllp* by P30 (Figure 3). Complementary ChIP–qPCR demonstrated increased H3K27me3 enrichment at promoter-proximal regions of *Sox10*, consistent with loss of KDM6B-mediated removal of Polycomb repressive marks (Figure 4). Together, these findings support a model in which KDM6B promotes oligodendrocyte development and postnatal myelination by maintaining *Sox10* in an active chromatin state.

Functionally, *Sox10* occupies a central position in the oligodendrocyte lineage hierarchy, acting upstream of key myelinogenic effectors such as *Mbp* and *Pllp* and being both necessary and sufficient to drive oligodendrocyte maturation(Stolt et al., 2002; Li et al., 2007; Pozniak et al., 2010). Several lines of evidence indicate that *Sox10* is a direct and functionally relevant KDM6B target *in vivo*: (i) *Sox10* is among the earliest and most robustly downregulated transcripts following *Kdm6b* deletion; (ii) the *Sox10* promoter gains repressive H3K27me3; and (iii) inducible *Sox10* expression partially rescues cortical myelination in *Kdm6b*-deficient cortices. The incomplete nature of this rescue suggests that additional KDM6B-dependent pathways—such as lipid/sterol biosynthetic programs or chromatin regulators that prime myelin gene loci—likely cooperate with SOX10 to achieve full oligodendrocyte maturation.

Our temporal manipulations further highlight the critical importance of developmental timing in lineage specification. Constitutive *Sox10* overexpression at E15.5 severely disrupted radial migration, differentiation, and proliferation of cortical progenitors, consistent with premature oligodendroglial commitment at the expense of normal neuronal differentiation. In contrast, delayed *Sox10* induction using a Tet-inducible system (E18.5–P16) permitted more appropriate radial migration into cortical layers and enabled partial rescue of cortical myelination (Figure 5). These findings underscore that epigenetic priming by KDM6B precedes and permits lineage consolidation, and that *Sox10* must be deployed within appropriate developmental windows to avoid perturbing cortical histogenesis.

An important aspect of our study is the Emx1-lineage focus. Although early cortical oligodendrocytes predominantly arise from ventral progenitors, dorsally derived Emx1 progenitors contribute substantially to postnatal oligodendrogenesis(Gorski et al., 2002; Tripathi et al., 2011; Crawford et al., 2016). By profiling Emx1-lineage nuclei, we demonstrate that KDM6B loss within this dorsal lineage is sufficient to suppress the *Sox10* → *Mbp* axis and impair cortical myelination, implicating KDM6B as a critical regulator of dorsally derived oligodendrocyte maturation.

Mechanistically, the concordance between epigenetic and transcriptional alterations strengthens causal inference. Accumulation of promoter-proximal H3K27me3 at *Sox10* is precisely the chromatin shift predicted upon loss of a H3K27me3 demethylase, and such modifications are tightly linked to transcriptional repression (Figure 3 and 4). These changes therefore likely represent direct, primary consequences of KDM6B loss.

Our findings also have clinical relevance. KDM6B variants are increasingly implicated in neurodevelopmental disorders, including autism spectrum disorder(Stessman et al., 2017; Stolerman et al., 2019; Satterstrom et al., 2020). Although neuroimaging phenotypes in patients are heterogeneous, our data predict that hypomorphic KDM6B states may compromise postnatal myelin maturation, potentially contributing to cognitive and behavioral deficits. The partial rescue achieved by *Sox10* suggests that targeting downstream nodes—such as SOX10-dependent transcriptional networks, lipid biosynthetic programs, or PRC2 antagonism—may represent tractable therapeutic strategies to restore myelin trajectories.

Several limitations warrant consideration. First, restriction to the Emx1 lineage complicates quantitative attribution across all oligodendrocyte sources; future fate-mapping and cell type–specific genetic studies will refine mechanistic resolution. Second, *Sox10* induction did not fully normalize myelination, which may reflect suboptimal timing or dosage, incomplete coverage of the SOX10 regulome, or contributions from *Sox10*-independent KDM6B targets. Third, we focused on promoter marks; enhancer regulation and three-dimensional chromatin architecture at myelin gene loci remain to be explored.

In summary, we propose that KDM6B licenses the oligodendrocyte differentiation program in the developing cortex by removing H3K27me3 and sustaining active chromatin at *Sox10* and its downstream myelin gene network. Loss of KDM6B blunts this program, delaying postnatal myelination, while temporally controlled *Sox10* induction partially compensates—highlighting SOX10 as a critical effector and revealing the multigenic nature of KDM6B-mediated myelin regulation. Future integration of cell type–specific genetics, single-cell multi-omics, and targeted epigenetic modulation will further elucidate how histone demethylation coordinates lineage maturation in the postnatal brain and how these pathways may be therapeutically leveraged in KDM6B-related disorders.

## Acknowledgements

We thank the MSU transgenic core facility to derive the *Kdm6b*-cKO founder mice. This work was supported by the National Institutes of Health (R01 GM127431, R01 MH130544).

## Author Contributions

J.H. conceived and supervised the project. Z.S. made the initial observations that led to project initiation, maintained the mouse colonies, and performed preliminary histological analyses. R.L. carried out all histological experiments and quantitative analyses. J.H. performed the RNA-seq and ChIP-qPCR experiments. G.M. and J.H. conducted the RNA-seq bioinformatic analyses. R.L., Z.S., and J.H. interpreted the data and wrote the manuscript.

