## supplemental materials for "Histone Demethylase KDM6B Promotes Postnatal Oligodendrocyte Development and Cortical Myelination"

.

**This PDF file includes:**

Figs. S1 to S3
Table S1

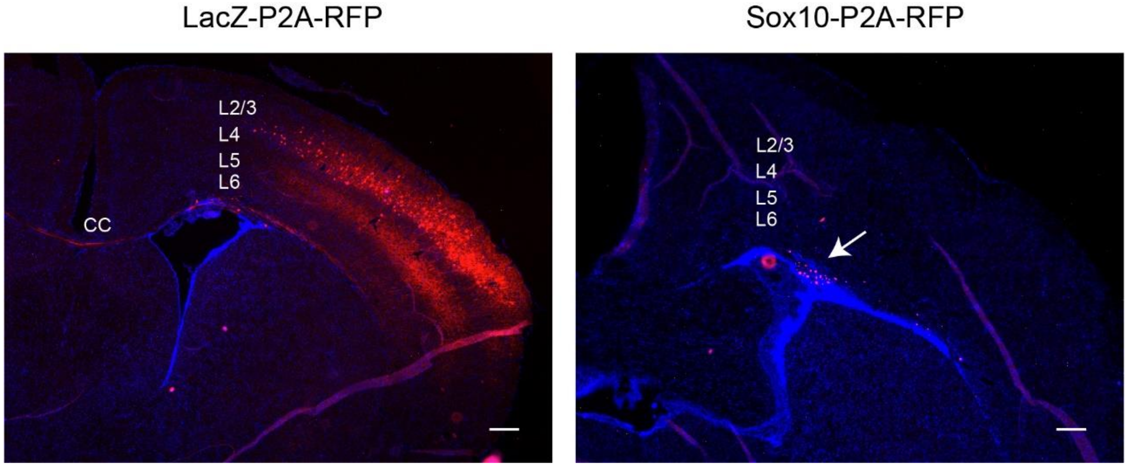


**Figure S1. Premature *Sox10* expression disrupts cortical progenitor migration, differentiation, and proliferation.**
Representative coronal cortical sections following in utero electroporation (IUE) at E15.5 with constitutively expressing LacZ–P2A–RFP (control) or Sox10–P2A–RFP constructs. In LacZ-electroporated cortices, $RFP^+$ cells migrate radially and populate upper cortical layers, exhibiting normal laminar distribution and cortical differentiation. In contrast, premature *Sox10* expression markedly disrupts radial migration, with the majority of $RFP^+$ cells accumulating in the subventricular zone adjacent to the corpus callosum (CC). *Sox10*-expressing cells also exhibit reduced dispersion and clustering near the ventricular region, consistent with impaired differentiation and proliferation of cortical progenitors. Nuclei are counterstained with DAPI (blue). Scale bars, 200 μm.

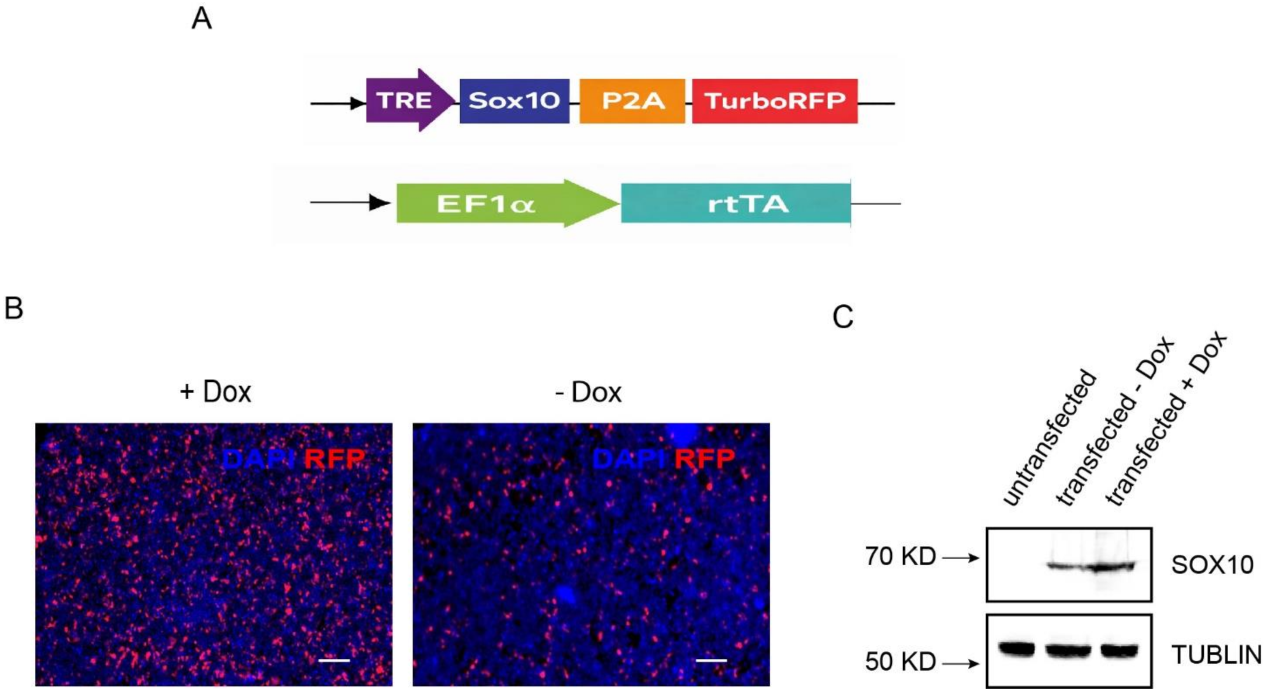


**Figure S2. Validation of doxycycline-inducible *Sox10* expression system.**
**(A)** Schematic of plasmids used for inducible Sox10 expression. The TRE promoter drives Sox10–P2A–RFP, allowing co-expression of SOX10 and RFP as separate proteins, while the EF1α promoter drives expression of rtTA for doxycycline (Dox)–dependent activation.
**(B)** Representative immunofluorescence images of HEK293T cells transfected with inducible Sox10 constructs in the presence (+Dox) or absence (−Dox) of doxycycline. Robust RFP expression is observed upon Dox treatment, with minimal basal expression in its absence. Nuclei are counterstained with DAPI (blue); RFP is shown in red. Scale bars, 20 μm.
**(C)** Immunoblot analysis of SOX10 protein levels in untransfected cells and transfected cells cultured with or without Dox. Dox treatment induces a marked increase in SOX10 expression, whereas little to no SOX10 is detected in the absence of Dox. Tubulin serves as a loading control.

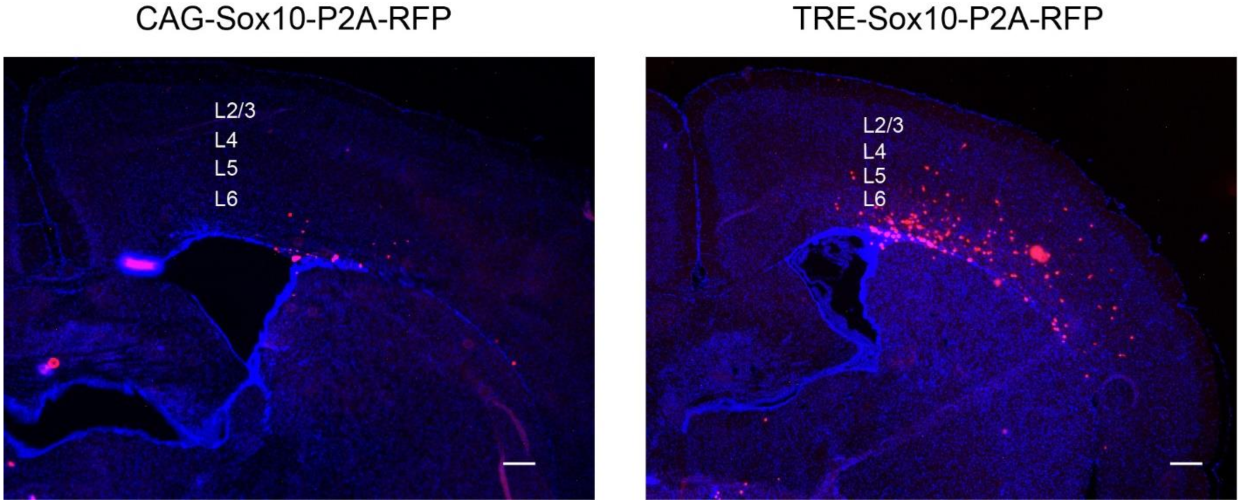


**Figure S3. Temporal control of *Sox10* expression improves radial migration, differentiation, and proliferation of cortical progenitors.**
Representative coronal cortical sections following in utero electroporation (IUE) at E15.5 with either constitutively active CAG–Sox10–P2A–RFP or doxycycline-inducible TRE–Sox10–P2A–RFP constructs. Constitutive *Sox10* expression driven by the CAG promoter severely restricts radial migration, with $RFP^+$ cells largely retained in periventricular regions. In contrast, temporally controlled *Sox10* induction using the TRE promoter permits radial migration of $RFP^+$ cells into upper cortical layers, consistent with improved differentiation and proliferation of cortical progenitors. Nuclei are counterstained with DAPI (blue). Scale bars, 200 μm.

**Supplementary Table 1. Sequences of all primers used in this study**

| Name | Sequence (5'-3') | Purpose |
|---|---|---|
| GAPDH-F | gcagtggcaaagtggagatt | qRT-PCR |
| GAPDH-R | gaatttgccgtgagtggagt | qRT-PCR |
| Kdm6b-F | agaggaaccagacagcactac | qRT-PCR |
| Kdm6b-R | cttccacctcttggcatca | qRT-PCR |
| ChIP_Sox10_amp1_F | ctgactgtgccacctgtatc | ChIP-qPCR |
| ChIP_Sox10_amp1_R | ctggactcagcttgggttt | ChIP-qPCR |
| ChIP_Sox10_amp2_F | aacgccttcatggtgtgg | ChIP-qPCR |
| ChIP_Sox10_amp2_R | cagagcttgcctagtgtcttg | ChIP-qPCR |
| ChIP_Sox10_amp3_F | ctgacccttgtagctccattt | ChIP-qPCR |
| ChIP_Sox10_amp3_R | cagcctaagatggttgtgatct | ChIP-qPCR |
| ChIP_Sox10_amp4_F | gcctgtctcttggtctcttac | ChIP-qPCR |
| ChIP_Sox10_amp4_R | ggaaagcgcctaaggaatct | ChIP-qPCR |
